# Rapid proteostasis controls monolayer integrity of quiescent endothelium

**DOI:** 10.1101/2022.08.09.503297

**Authors:** Fabienne Podieh, Roos Wensveen, Max C. Overboom, Lotte Abbas, Jisca Majolée, Peter L. Hordijk

## Abstract

Endothelial monolayer permeability is regulated by actin dynamics and vesicular traffic. Recently, ubiquitination was also implicated in the integrity of quiescent endothelium, as it differentially controls the localization and stability of adhesion- and signaling proteins. We found that inhibition of E1 ubiquitin ligases induces a rapid, reversible loss of integrity in quiescent, primary human endothelial monolayers, accompanied by increased F-actin stress fibers and the formation of intercellular gaps. Concomitantly, total protein and activity of the actin-regulating GTPase RhoB, but not its close homologue RhoA, increase ∼10-fold in 5-8 h. The depletion of RhoB, but not of RhoA, the inhibition of actin contractility and the inhibition of protein synthesis all significantly rescue the loss of cell-cell contact induced by E1 ligase inhibition. Our data suggest that in quiescent human endothelial cells, the continuous and fast turnover of short-lived proteins that negatively regulate cell-cell contact, is essential to preserve monolayer integrity.

## Introduction

The most inner lining of blood vessels is formed by the endothelium which acts as a dynamic barrier controlling the extravasation of leukocytes and plasma from the circulation into the tissue. Dysfunction of the endothelial barrier leads to vascular leak and edema and is a hallmark of chronic inflammatory diseases. Thus, control of endothelial integrity is crucial for tissue and organ health. This integrity is critically dependent on cell-cell adhesion between endothelial cells (ECs), which is mediated by the homotypic adhesion protein vascular endothelial (VE)-cadherin. Intracellular adaptor proteins link VE-cadherin to the cortical actin cytoskeleton, stabilizing its adhesive function (Perez-Moreno et al. 2006). Importantly, both this connection and the organization of the actin cytoskeleton are dynamic and subject to complex regulation. For example, agonist-induced changes in cytoskeletal organization in combination with the induction of myosin-based contraction can generate contractile forces that disrupt VE-cadherin-based complexes and impair vascular integrity (Giannotta, Trani, and Dejana 2013; Hordijk et al. 1999; Vandenbroucke et al. 2008).

The key regulators of cytoskeletal dynamics in ECs are members of the family of Rho GTPases. RhoA/B activity promotes F-actin stress fiber assembly and myosin activation leading to cell contraction, loss of cell-cell contact, and an increase in vascular permeability. In contrast, Rac1 promotes actin polymerization and the formation of a cortical F-actin network improving endothelial integrity (Pronk et al. 2019; Wojciak-Stothard and Ridley 2002). Signaling by Rho GTPases is primarily controlled by their cycling between an inactive, GDP-bound state and an active, GTP-bound state. This cycle is tightly regulated both in time and space by activating guanine nucleotide exchange factors (GEFs) and inactivating GTPase-activating proteins (GAPs) (Cherfils and Zeghouf 2013; Bos, Rehmann, and Wittinghofer 2007).

In addition to their GTP/GDP cycling, Rho GTPases can be post-translationally modified by lipidation, phosphorylation, sumoylation or ubiquitination to further regulate their localized signaling capacity. Ubiquitination entails the covalent attachment of the highly conserved, 76 amino acid ubiquitin peptide to lysine residues of a substrate and subsequently to lysine residues of ubiquitin itself (known as poly-ubiquitination). The formation of such chains through ubiquitin K48 or K63 residues, targets the substrate for proteasomal or lysosomal degradation, respectively (Akutsu, Dikic, and Bremm 2016; Komander and Rape 2012). Mechanistically, E1, E2 and E3 ubiquitin ligases act in a cascade, with the E3 ligase transferring the activated ubiquitin to the substrate (Pickart and Eddins 2004). Rho GTPases are targeted by several ubiquitin E3 ligases. For example, HACE1 and IAP target Rac1 for degradation, thereby controlling the generation of reactive oxygen species (ROS) in HUVEC as well as cell spreading and migration in HEK293 and HeLa cells (Torrino et al. 2011; Daugaard et al. 2013; Oberoi et al. 2012). RhoA stability is regulated by the Smurf1 and Cullin3-BACURD E3 ligases to control cell polarity and cell migration in tumor- and HeLa cells, respectively (Wang et al. 2003; Chen et al. 2009). In addition, Rho targeting by Cullin3 limits smooth muscle cell contractility and hypertension (Pelham et al. 2012; Agbor et al. 2016)

The ubiquitously expressed E1 ligases ubiquitin-activating enzyme 1 (UBA1) and UBA6 initiate the ubiquitination cascade by activating ubiquitin and charging the E2 ligases (Schulman and Harper 2009). Recently, Hyer et al. identified a small molecule inhibitor named MLN7243 (a.k.a. TAK-243) targeting both UBA1 and UBA6, thus inhibiting cellular ubiquitination (Hyer et al. 2018). Prolonged inhibition of the E1 enzymes leads to impairment of DNA damage repair, cell cycle progression and proliferation and exhibits antitumor activity in various tumor cell lines and in mouse xenograft models for solid and hematological cancers (Hyer et al. 2018; Barghout et al. 2019; Zhuang et al. 2019; Liu et al. 2020; Best et al. 2019). The therapeutic potential of MLN7243 is currently being assessed in patients with advanced solid tumors or leukemia (ClinicalTrials.gov Identifier: NCT02045095 and NCT03816319).

We previously identified RhoB, rather than RhoA, as a major negative regulator of monolayer integrity in quiescent endothelial cells (Amado-Azevedo et al. 2017; Pronk et al. 2019). RhoB has a very short half-life (1-2 h), as compared to RhoA (8-12 h), and we showed that ubiquitination of RhoB by the Cul3-Rbx1-KCTD10 E3 ligase is a key regulatory event controlling endothelial cell-cell contact (Kovacevic et al. 2018). However, whether there exists a more generic role for fast protein turnover in regulating endothelial integrity is unknown. To address this, we used short-term inhibition of the E1 ubiquitin ligases in quiescent, primary human endothelial cells and analyzed the consequences for barrier integrity as well as for the turnover of RhoA, RhoB and Rac1. Our results show that E1 inhibition rapidly (within 5-8 h) and reversibly induces marked cytoskeletal changes and a pronounced loss of endothelial integrity. This is accompanied by a fast accumulation of total and activated RhoB, but not RhoA or Rac1. Finally, this effect is completely dependent on protein synthesis. Together, these results support the concept that continuous and rapid protein turnover is required to preserve the integrity of quiescent endothelium.

## Results

### Ubiquitination preserves endothelial barrier function

To establish the importance of ubiquitination for endothelial barrier integrity, the ubiquitination pathway in primary HUVEC monolayers was inhibited by the E1 ligase inhibitor MLN7243 (TAK-243; Figure 1A). MLN7243 caused a rapid, dose-dependent reduction of endothelial integrity within 4-8 h, as analyzed in real time using ECIS (Figure 1B). Additionally, increased permeability of endothelial monolayers was observed after treatment with MLN7243, as quantified by HRP leakage in a transwell assay (Figure 1C). The total pool of ubiquitinated proteins decreased accordingly in a dose- and time-dependent manner (Figure S1A). Since ubiquitination plays a crucial, regulatory role in many signaling events, inhibition of the E1 ligase may eventually compromise cell viability. To address this, we studied time-dependent changes in caspases, marking the induction of apoptosis. MLN7243 treatment of HUVEC resulted in a reduction of caspase 9 and an increase in cleaved caspase 3 as detected after 16 h, but not during the first 8 h of incubation (Figure S1B), indicating that the MLN7243-induced loss of monolayer integrity was not due to apoptosis.

**Figure 1.**
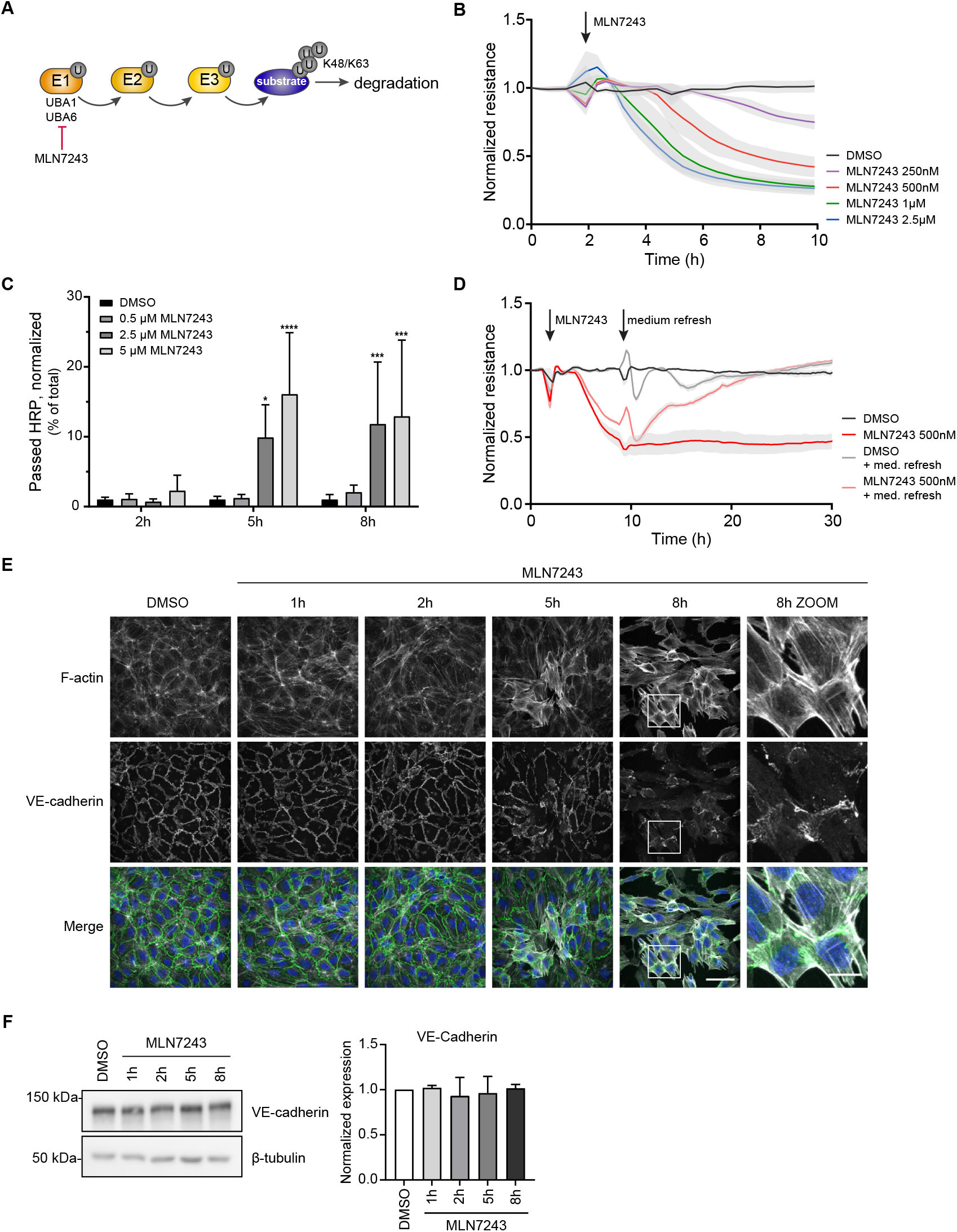
MLN7243 disrupts endothelial barrier integrity. (**A**) MLN7243 inhibits the E1 ligases UBA1 and UBA6. (**B**) Normalized resistance of HUVEC treated with different concentrations of MLN7243. (**C**) Macromolecule passage of HRP across HUVEC monolayer treated with 500 nM MLN7243, normalized to their respective DMSO control. (data are presented as mean + SD, n=3) *, p<0.05, ***, p<0.001, ****, p<0.0001 (**D**) Normalized resistance of HUVEC treated with 500 nM MLN7243 and refreshed with medium without MLN7243 after 7 h of treatment. (**E**) Monolayer of HUVEC treated with 500 nM MLN7243 for the indicated times were stained for F-actin (white) and VE-cadherin (green) and counterstained with DAPI (blue). Scale bars represent 50 μm in overview images and 15 μm in zoomed images. (**F**) Western blot analysis of VE-cadherin expression after 500 nM MLN7243 treatment for the indicated times. β-tubulin was used as loading control. Bar graph shows quantification of VE-cadherin expression normalized to β-tubulin (data are presented as mean + SD, n=3). See also figure S1.

To further exclude that the MLN7243-induced loss of integrity was the result of irreparable cell damage, we tested the reversibility of its effect. We found that the reduction of transendothelial resistance, induced by MLN7243, was fully restored when the compound was removed after 7 h by washout. This induced a complete recovery of endothelial electrical resistance within 10 h (Figure 1D), indicating that reformation of cell-cell contact, rather than increased cell proliferation, underlies the restoration of barrier function. MLN7243 did not induce an upregulation of ICAM1, suggesting that E1 inhibition does not trigger an inflammatory response, which is generally accompanied by a loss of barrier function (Figure S1C). Similar as for HUVEC, inhibiting ubiquitination with MLN7243 in hMVEC resulted in a dose-dependent impairment of endothelial integrity (Figure S1D). Based on these initial findings, we decided for subsequent experiments to limit both the dose (500 nM) and duration (max. 8 h) of treatment with MLN7243.

To further understand the barrier-disruptive effect of E1 inhibition, distribution of VE-cadherin and F-actin was analyzed by immunofluorescence microscopy. At early time points of incubation with MLN7243 (1 h and 2 h), cell-cell junctions appeared stable as indicated by honeycomb-like structures marked by VE-cadherin and cortical F-actin. However, after 5 h and 8 h of exposure to MLN7243, F-actin stress fibers were strongly increased and VE-cadherin localization at junctions was markedly reduced (Figure 1E, S1E). The levels of VE-cadherin protein remained unaltered (Figure 1F), suggesting an indirect effect of MLN7243 on VE-cadherin distribution. Together, these findings show that short term inhibition of ubiquitination in EC has inhibitory, but reversible effects on VE-cadherin-mediated cell-cell contact. This shows that in quiescent EC, there is continuous ubiquitin-mediated degradation of barrier-disrupting protein(s). Moreover, these data further identify in primary human EC, a dynamic proteostatic mechanism that acts at short time scales and preserves monolayer integrity.

### MLN7243 causes accumulation of active RhoB, but not RhoA

Rho GTPases are key regulators of actin filament organization. RhoA/B/C, which are identical in their effector domain (Schaefer et al., 2014), promote formation of contractile F-actin fibers via their shared effector Rho-associated kinase (ROCK). ROCK, in turn, regulates phosphorylation of Myosin Light Chain (MLC) and the induction of actomyosin contraction (Komarova et al. 2017). The increased F-actin stress fiber formation which we observed following E1 inhibition in HUVEC with MLN7243 (Figure 1E) was accompanied by elevated levels of phosphorylated MLC (Figure 2A). To test if cell contraction upon E1 inhibition is mediated by ROCK, HUVEC were pre-treated with the ROCK inhibitor Y27632. This significantly reduced MLN7243-induced disruption of endothelial integrity, indicative for a role of ROCK-mediated contraction (Figure 2B, C).

**Figure 2.**
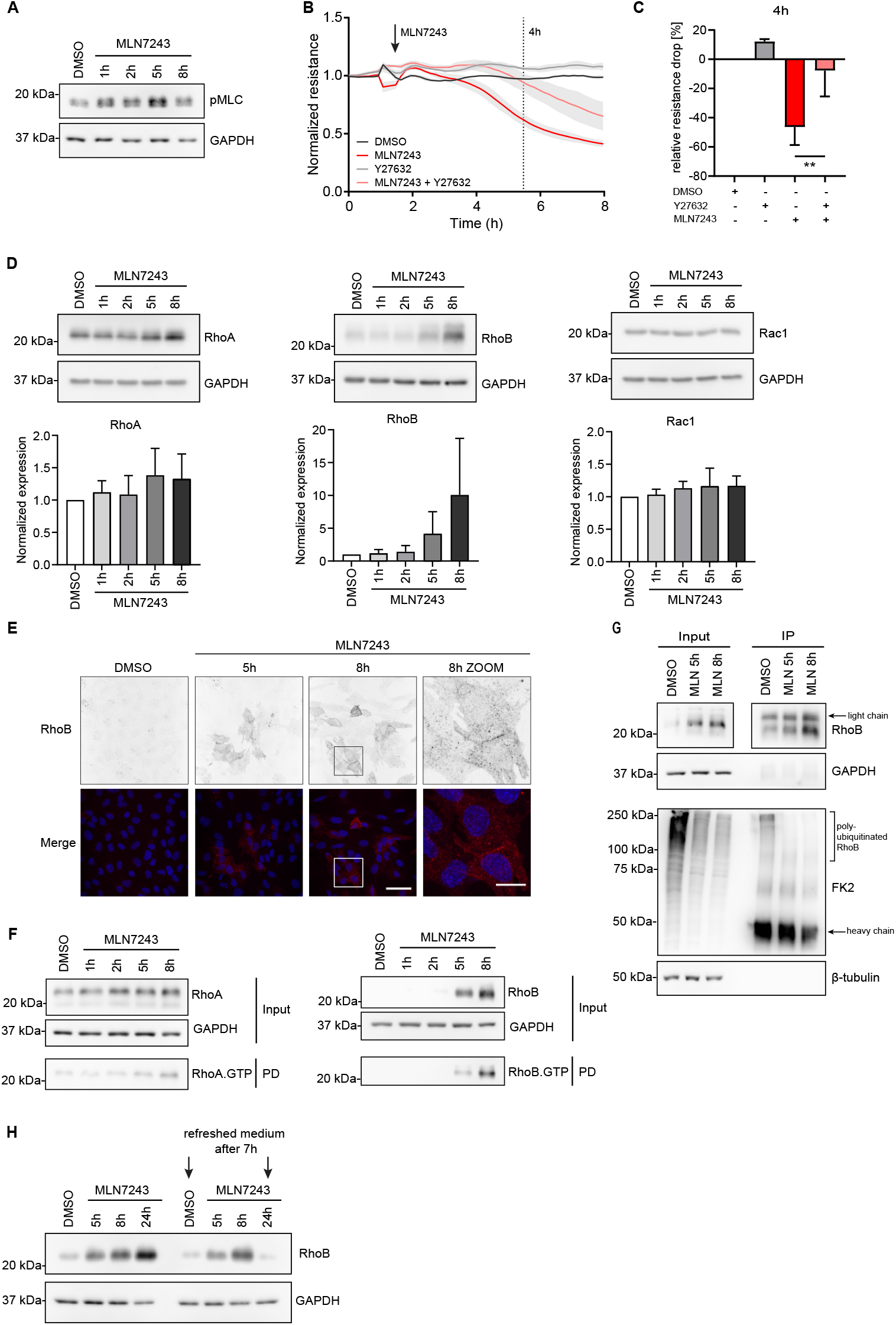
Inhibition of ubiquitination by MLN7243 leads to increased RhoB expression and activity. (**A**) Western blot analysis for pMLC of HUVEC treated with 500 nM MLN7243 for the indicated times. GAPDH was used as loading control. (**B, C**) Normalized resistance of HUVEC treated with 500 nM MLN7243, 10 μM Y27632 or both, and (**C**) relative drop in resistance after 4h (data presented as mean + SD, n=3). **, p<0.01 (**D**) Western blot analysis and quantification of RhoA, RhoB and Rac1 protein expression in HUVEC treated with 500 nM MLN7243. Bar graph shows quantification of RhoA/B, Rac1 expression, relative to GAPDH and DMSO (data are presented as mean + SD, n=3). (**E**) Immunofluorescent staining for RhoB (red) and counterstained with DAPI (blue) of HUVEC after treatment with 500 nM MLN7243. Scale bars represent 50 μm in overview images and 15 μm in zoomed images. (**F**) Rhotekin pulldown of HUVEC treated with 500 nM MLN7243 followed by Western blot analysis for the GTP-bound, active forms of RhoA and RhoB. GAPDH was used as loading control. (**G**) Endogenous RhoB was immunoprecipitated from HUVEC treated with 500 nM MLN7243 followed by Western blot analysis of RhoB and ubiquitinated proteins (FK2). Input equals 8%. (**H**) Western blot analysis of RhoB expression after treatment with 500 nM MLN7243 for the indicated times and with a medium refresh without MLN7243 after t = 7 h (right panel only). See also figure S2.

As ROCK acts downstream of Rho GTPases, we investigated if the short-term inhibition of ubiquitination alters the levels and activity of the main barrier regulating Rho GTPases RhoA, RhoB and Rac1. Strikingly, MLN7243 induced a 5-10-fold increase in RhoB protein level within 5-8 h, whereas the levels of RhoA and Rac1 showed only little (∼1.4 fold for RhoA) to no increase (Rac1) (Figure 2D). Similar to the MLN7243-induced loss of endothelial integrity, the MLN7243-induced increase in RhoB level was dose-dependent (Figure S2A). Similar as in HUVEC, MLN7243 caused a strong accumulation of RhoB in hMVEC, whereas RhoA was only slightly increased (Figure S2B). In line with these findings, immunofluorescence microscopy revealed a clear increase in RhoB levels after 5 h and 8 h of exposure of HUVEC to MLN7243 (Figure 2E). Under these conditions, RhoB localizes to intracellular vesicles throughout the cytosol and close to, or at cell-cell contacts. In parallel, RhoB activity, analyzed by Rhotekin pulldown, was markedly increased after 5 h and 8 h of MLN7243 treatment (Figure 2F). This was in marked contrast to the effect on RhoA which showed only a minor increase in activity (Figure 2F). The loss of RhoB ubiquitination induced by MLN7243 was further confirmed by an IP of endogenous RhoB in HUVEC. After 5 h and 8 h of MLN7243 treatment, the amount of poly-ubiquitinated RhoB was clearly diminished (Figure 2G). Finally, similar to the loss of integrity, the MLN7243-induced increase in RhoB was reversible since washout of MLN7243 after 7 h caused a strong reduction of RhoB protein level, comparable to control levels (Figure 2H). This data shows that continuous ubiquitination plays a crucial role in limiting the levels of endothelial RhoB protein and activity, with only minor effects on RhoA.

In various cancer cell lines, the inhibition of the E1 ligases results in the accumulation of myriad proteins and is associated with the induction of ER stress (Hyer et al. 2018; Best et al. 2019; Liu et al. 2021). In line with this, MLN7243 induced ER stress in HUVEC after 8 h as indicated by elevated phosphorylation of eIF2α (Figure S2C). Induction of ER stress by Tunicamycin (TM) resulted in a rapid disruption of the endothelial integrity, comparable to the effects of MLN7243 (Figure S2D, E). While this is in line with the notion that ER-stress impairs endothelial integrity (Finnie and O’Shea 1990; Lenin et al. 2019), TM treatment did not increase RhoB levels (Figure S2D). This indicates that the observed increase in RhoB protein, induced by MLN7243, is consequent to the inhibition of ubiquitination rather than to ER-stress.

### Ubiquitination of RhoB, but not RhoA, is crucial for endothelial barrier function

Based on the strong upregulation of RhoB following E1 inhibition, we next analyzed the requirement for RhoB expression in the inhibitory effects of MLN7243 on endothelial integrity. Following the siRNA-induced loss of RhoB and RhoA expression, HUVEC were treated with MLN7243 and barrier function was measured by ECIS. Interestingly, silencing of RhoB partly rescued the loss of barrier function induced by MLN7243, whereas depletion of RhoA did not (Figure 3A, B). Importantly, siRNA-mediated loss of RhoB induces an increase of RhoA protein level and vice versa (Figure 3C) (Pronk et al. 2019). Since this might partially compensate for the loss of RhoB or RhoA function, confounding the interpretation of this experiment, we also tested the effects of simultaneous depletion of RhoA and RhoB. Combined knockdown of these two Rho GTPases resulted in a pronounced and significant protection of endothelial barrier integrity (Figure 3D, E), largely preventing the effect of MLN7243. This indicates that lack of, primarily, RhoB makes the endothelial barrier refractory to the short-term effects caused by inhibition of cellular ubiquitination. These data support the notion that continuous ubiquitination and degradation of RhoB is crucial to preserve barrier function in quiescent endothelial cells.

**Figure 3.**
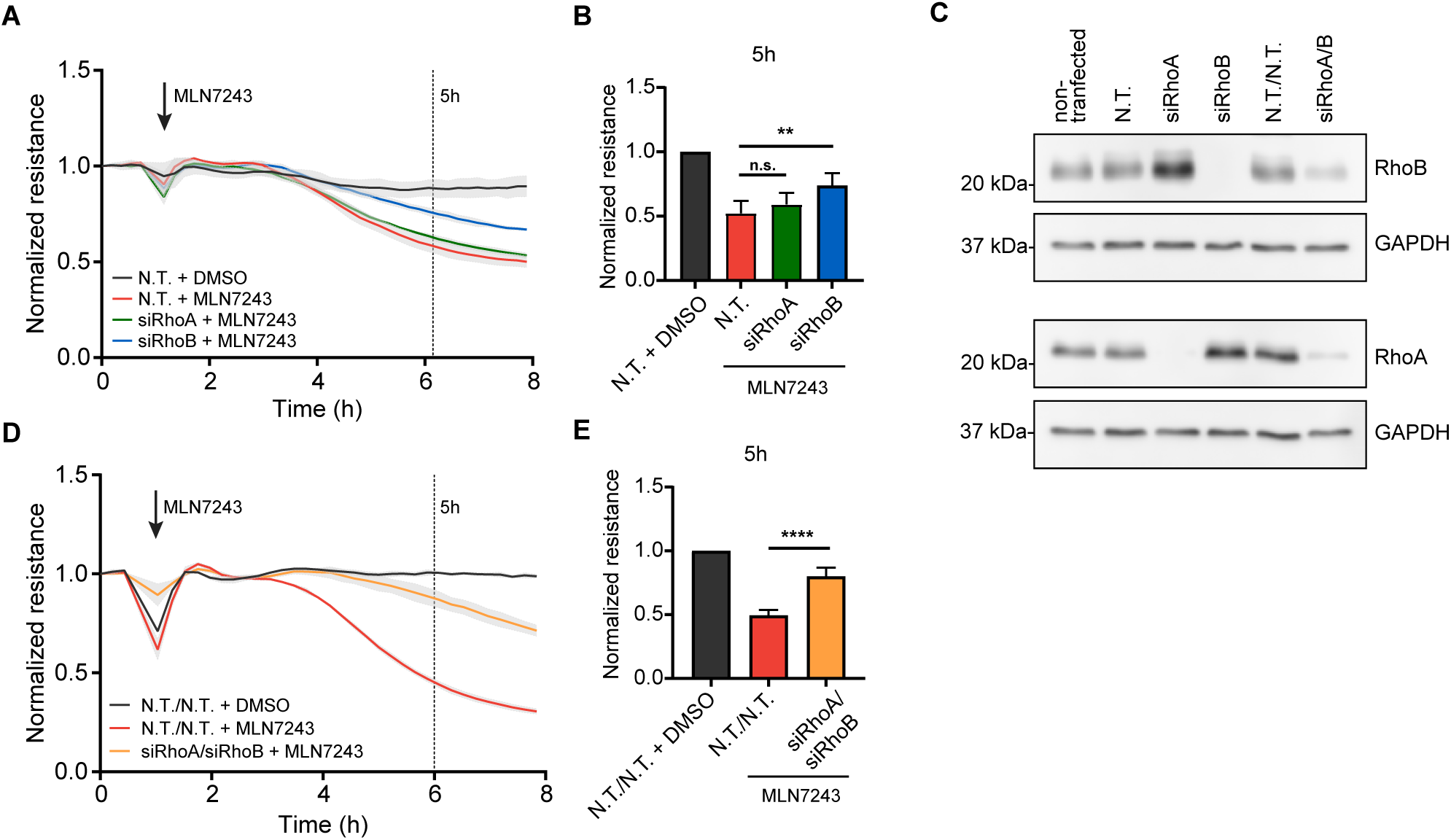
Depletion of RhoB partially rescues the MLN7243-induced loss of endothelial integrity. (**A-E**) HUVEC were transfected with non-targeting control siRNA (N.T.), siRhoA and/or siRhoB and cultured for 72 h. (**A, B**) Normalized resistance of HUVEC with single knockdowns treated with 500 nM MLN7243 (**B**) Bar graphs represent mean + SD of normalized endothelial resistance after 5 h of four individual experiments relative to DMSO. **, p<0.01. (**C**) Western blot analysis of RhoB and RhoA expression. (**D, E**) Normalized endothelial resistance of HUVEC with double knockdowns of RhoA/B treated with 500 nM MLN7243. (**E**) Bar graphs represent mean + SD of normalized endothelial resistance after 5 h, relative to the DMSO control, of four individual experiments. ****, p<0.0001.

### Inhibition of protein translation prevents the loss of endothelial barrier function induced by MLN7243 and MLN4924

The rapid effects of MLN7243 on barrier integrity suggest that dynamic proteostasis controls endothelial integrity in quiescent monolayers. However, protein turnover is a function of both ubiquitin-mediated degradation and protein synthesis. To address the contribution of protein synthesis to this effect, HUVEC were briefly (2 h) pre-treated with the protein translation inhibitor cycloheximide (CHX) and barrier function was measured with ECIS. Intriguingly, CHX by itself caused a marked improvement of endothelial integrity within 6-8 h (Figure 4A, B), indicating that in quiescent EC, barrier-disrupting proteins with a very short half-life are continuously synthesized. In line with this notion, CHX completely prevented the barrier-disruptive effect of MLN7243 (Figure 4A, B). Since RhoB is known to have a short half-life (1-2 h) (Figure S3A, (Lebowitz, Davide, and Prendergast 1995)), we tested if CHX affects the MLN7243-induced accumulation of RhoB. Indeed, inhibition of translation completely blocked the increase in RhoB, whereas the levels of RhoA and Rac1 (both with longer half-lives; Figure S3A) remained unaffected (Figure 4C). Concomitantly, CHX prevented both the loss of junctional localization of VE-cadherin and formation of F-actin stress fibers, induced by E1 inhibition (Figure 4G). Importantly, CHX did not alter VE-cadherin levels or junctional distribution (Figure S3B, 4G) or Rac1 levels or activity within the time frame of the experiment (Figure S3C). These data suggest that the ongoing synthesis of signaling-competent RhoB plays a prominent role in actomyosin contractility and (local) barrier instability, which is kept in check by its fast and continuous ubiquitination and degradation.

**Figure 4.**
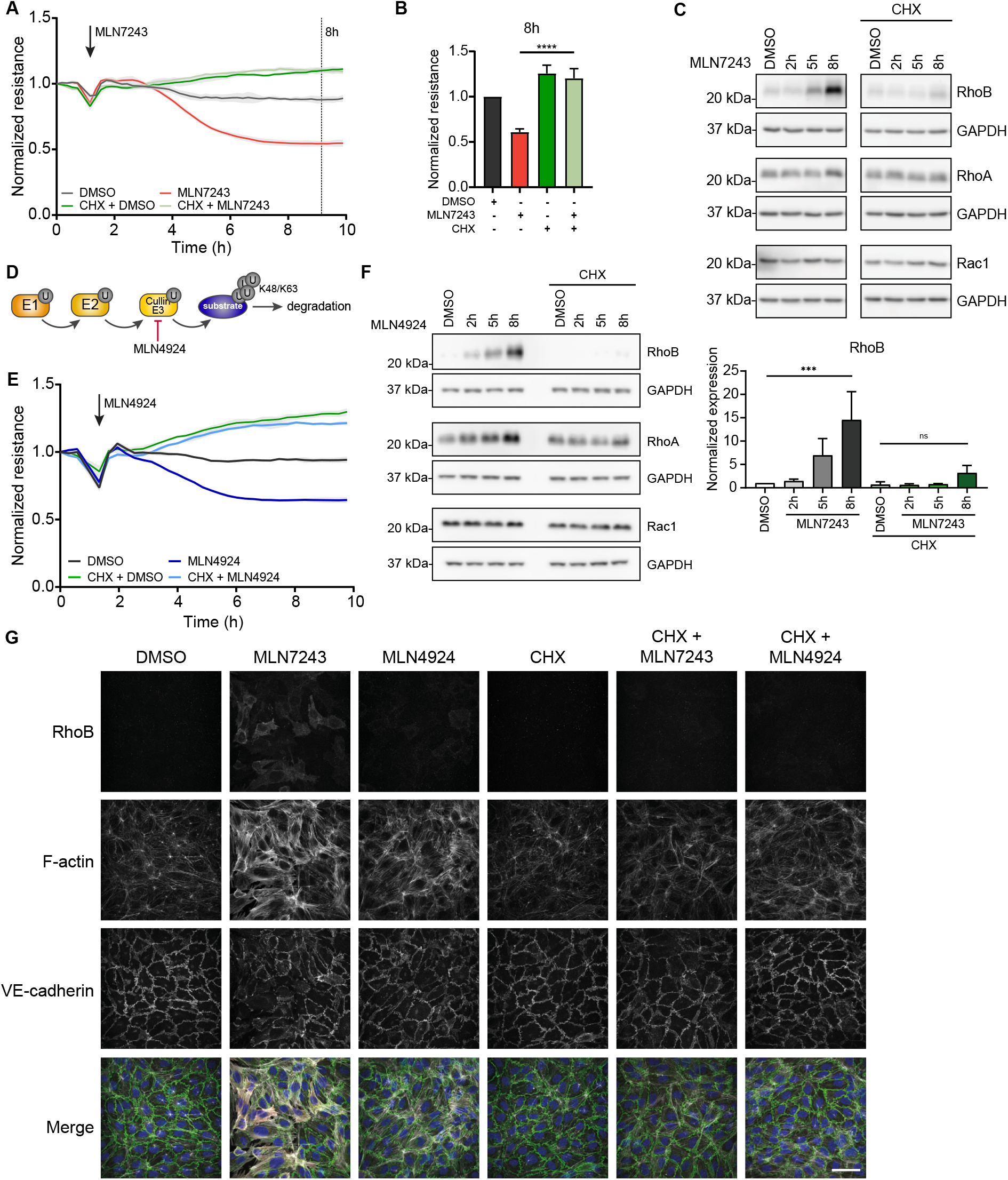
Regulation of RhoB turnover is crucial for endothelial integrity. (**A-C**) HUVEC were treated with 500 nM MLN7243, following pre-treatment as indicated for 2 h with 0.5 μg/ml CHX to block protein synthesis. (**A, B**) Normalized resistance (**B**) Bar graphs represent mean + SD of normalized endothelial resistance after 8 h, relative to DMSO (n=3). ****, p<0.0001. (**C**) Western blot analysis for RhoB, RhoA and Rac1 expression. GAPDH is used as loading control. Quantification of RhoB expression represents mean + SD, relative to GAPDH and DMSO control. n=3, ***, p<0.001 (**D**) MLN4924 inhibits cullin-based E3 ligases. (**E**) Normalized resistance and (**F**) Western blot analysis for expression of RhoB, RhoA and Rac1 after treatment of HUVEC with 500 nM MLN4924 and pre-treatment for 2 h with 0.5 μg/ml CHX. GAPDH was used as loading control. (**G**) Immunofluorescent staining for RhoB (red), F-actin (white) and VE-cadherin (green) and counterstained with DAPI (blue) of HUVEC after pre-treatment of 0.5 μg/ml CHX for 2 h followed by 5 h MLN7243 or MLN4924. Scale bar represents 50 μm. See also figure S3.

Previously, we showed that inhibition of Cullin-based E3 ligases by the neddylation inhibitor MLN4924 leads to an accumulation of RhoB and a concomitant loss of endothelial barrier function (Figure 4D; (Kovacevic et al. 2018). Thus, we investigated if these MLN4924-induced effects are also dependent on protein synthesis. In accordance with the above, CHX completely blocked barrier breakdown by Cullin inhibition and prevented the concomitant increase in RhoB protein level (Figure 4E, F). Consistently, inhibition of protein synthesis also prevented the stress fiber formation and cortical actin bundles induced by MLN4924 (Figure 4G). This further confirms the notion that RhoB synthesis and its Cullin3-mediated degradation is tightly regulated to preserve endothelial integrity.

### Regulation of RhoB synthesis in inflamed endothelium

While the above experiments are all performed in quiescent endothelial monolayers, various receptor agonists are known to activate ECs and disrupt monolayer integrity within a ∼4-8 h time frame. The pro-inflammatory cytokine TNFα is known to enhance vascular permeability and to promote transcription of RhoB mRNA leading to elevated RhoB protein level and activity, all within 4-8 h (Kroon et al. 2013). In accordance with these findings, both the TNFα-induced increase in RhoB protein and the concomitant loss of endothelial integrity were prevented by CHX (Figure 5A, B). Similarly, the pro-inflammatory cytokine interleukin-1β (IL-1β) induced accumulation of RhoB and disruption of barrier function, which was both prevented by CHX (Figure 5C, D). This suggests that tight regulation of RhoB turnover is crucial in not only in quiescent but also in inflamed endothelium.

**Figure 5.**
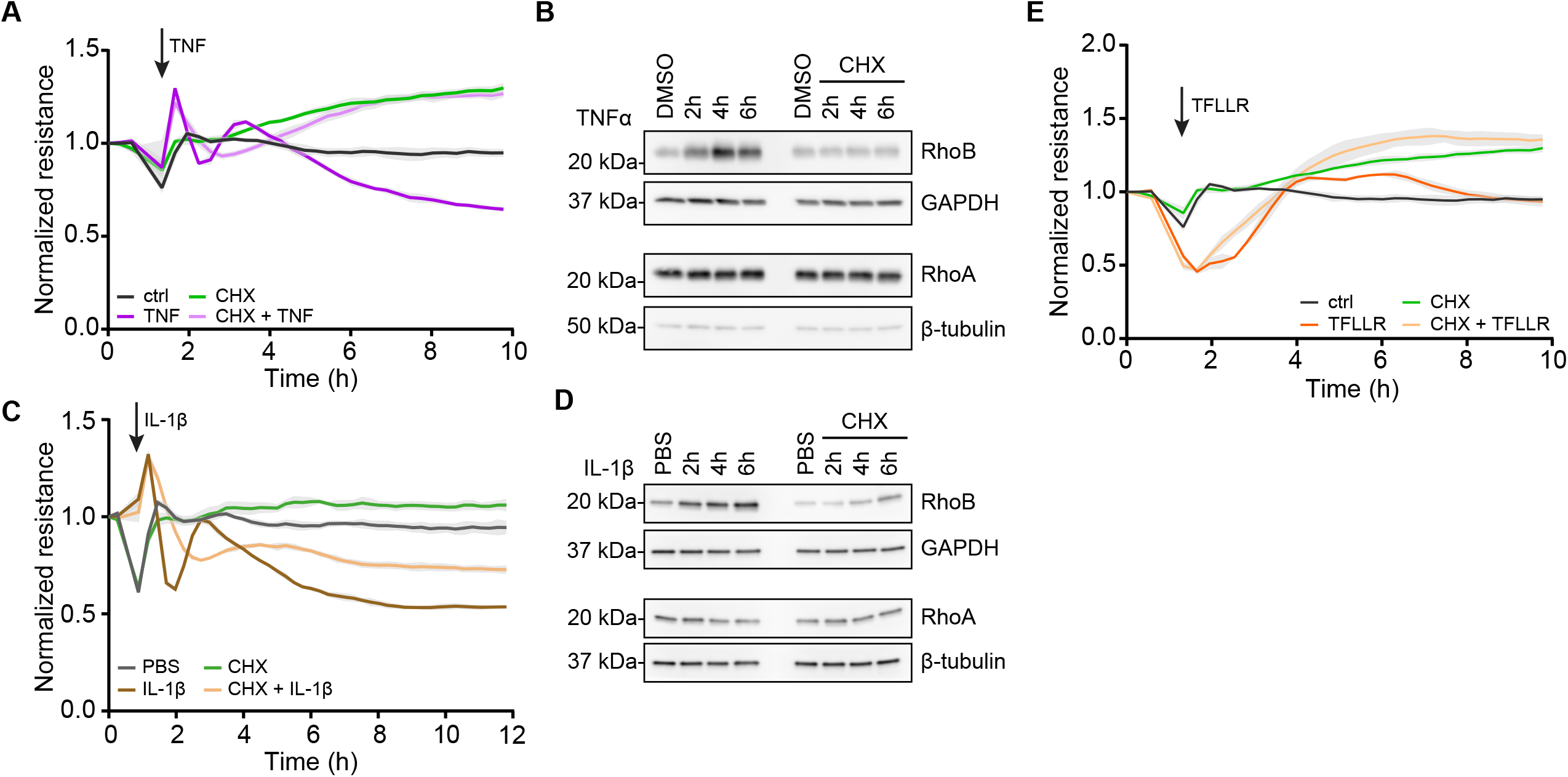
CHX prevents loss of endothelial integrity induced by pro-inflammatory cytokines. (**A**) Normalized resistance and (**B**) Western blot analysis for expression of RhoB and RhoA expression after HUVEC were pre-treated with 0.5 μg/ml CHX for 2 h and treated with 10 ng/ml TNFα. GAPDH and β-tubulin were used as loading controls. (**C**) Normalized resistance of HUVEC after pre-treatment with 0.5 μg/ml CHX for 2 h and treatment with 100 μM TFFLR. (**D**) Normalized resistance and (**E**) Western blot analysis for RhoA and RhoB expression after HUVEC were pre-treated with 0.5 μg/ml CHX for 2 h and treated with 10 ng/ml IL-1β for indicated times. GAPDH was used as loading control.

In addition to TNFα, activation of the G-protein coupled protease activated receptor 1 (PAR1) by thrombin or the Thrombin Receptor Peptide (TRP, (Minnear et al. 1993)) is a widely used approach to investigate the mechanisms underlying endothelial integrity. Activation of PAR1 induces a very fast and transient loss of endothelial barrier function, which is accompanied by, and dependent on, a pronounced activation of RhoA (van Nieuw Amerongen et al. 2000). Interestingly, inhibition of protein synthesis did not affect the TRP-induced transient loss of barrier function (Figure 5E). Together, these experiments show that the relatively slow and persistent induction of barrier loss induced by TNFα and IL-1β is mediated by (upregulation of) RhoB, while the fast and transient effect induced by PAR1 is mediated by the activation of RhoA and independent of protein translation.

## Discussion

In this study, we report that inhibiting cellular ubiquitination by the E1 ligase blocker MLN7243 (a.k.a. TAK-243) in primary human venous and microvascular EC causes a marked increase in densely packed F-actin stress fibers and an increase in phosphorylation of MLC. The resulting contraction is accompanied by a redistribution of VE-cadherin away from cell-cell junctions and a loss of cell-cell contact. These effects occur already within 5-8 h, are fully reversible and cannot be explained by ER stress or apoptosis. In contrast, they are completely dependent on protein synthesis. Together, these data show that the continuous synthesis and ubiquitin-mediated degradation of proteins with a short half-life plays a central role in the integrity of quiescent endothelial monolayers.

We recently identified Cullin3-Rbx1-KCTD10 as the E3 ligase targeting RhoB for K63 poly-ubiquitination and lysosomal degradation in quiescent human ECs (Kovacevic et al. 2018). Similarly, the Cullin3-KCTD10 E3 ligase has been recently shown to target RhoB for degradation in HER2-positive breast cancer cells (Murakami et al. 2019). In marked contrast to RhoA and Rac1, RhoB has a very short half-life of 1-2 h (Lebowitz, Davide, and Prendergast 1995). This is likely due to the fact that RhoB does not bind the ubiquitously expressed chaperone RhoGDI1, which is known to protect GTPases from activation and degradation (Garcia-Mata, Boulter, and Burridge 2011; Michaelson et al. 2001; Mosaddeghzadeh et al. 2021; Boulter et al. 2010). Consequently and because of the abundance of GTP in the cytosol, newly synthesized RhoB rapidly becomes GTP-bound, signaling competent and sensitive to degradation. Consistent with this idea, our data suggest that the regulation of RhoB output occurs predominantly at the level of its expression and degradation. We show that inhibiting cellular ubiquitination, i.e. blocking the E1 ubiquitin ligase by MLN7243, leads to a rapid, ≥10-fold increase in total RhoB protein level and activity, coinciding with the induction of contraction and disruption of the endothelial barrier. Notably, silencing of RhoB, but not of RhoA, significantly rescues the MLN7243-induced loss of barrier function. These data support the idea that continuous ubiquitin-mediated degradation of RhoB is essential to preserve quiescent endothelial integrity.

RhoB and RhoA share 88% sequence homology (Schaefer et al., 2014) and are both well-known negative regulators of endothelial barrier function by inducing ROCK-mediated F-actin contraction (Wojciak-Stothard and Ridley 2002). Ubiquitination of active and inactive RhoA by various E3 ligases and in various different cell types has been described previously. Smurf1, SCF(FBXL19) and Cullin3-BACURD were identified as E3 ligases that mediate ubiquitination of (active and inactive) RhoA, leading to its degradation. This mechanism contributes to the regulation of tumor cell motility, neurite outgrowth, proliferation and blood pressure (Wang et al. 2003; Bryan et al. 2005; Wei et al. 2013; Chen et al. 2009; Pelham et al. 2012). Interestingly, in ECs, the inhibitors of apoptosis proteins (IAPs) were suggested to modulate RhoA activity and, thereby, endothelial barrier function in response to thrombin (Hornburger et al. 2014). In contrast, our data show that in quiescent EC, RhoA ubiquitination does not significantly control barrier function as E1 ligase inhibition induces only a small increase in RhoA abundance and activity. Additionally, depletion of RhoA only marginally improves endothelial resistance when the ubiquitination cascade is inhibited, This indicates that ubiquitin-mediated degradation of RhoA plays a minor role in the integrity of quiescent endothelial monolayers.

Protein turnover is not only governed by the rate of ubiquitin-dependent degradation but also by protein synthesis. Intriguingly, inhibition of protein synthesis is sufficient to enhance quiescent endothelial integrity within 4 h and to completely prevent MLN7243-induced cytoskeletal changes, and loss of junctional VE-cadherin localization and monolayer integrity. These data reveal that in quiescent EC, barrier function is controlled by negative regulators with a short half-life. In addition, MLN7243-induced accumulation of RhoB is abolished by CHX. In good agreement with these results, barrier disruption and increase in RhoB protein level, induced by the Cullin E3 ligase inhibitor MLN4924 (Kovacevic et al. 2018), also requires protein synthesis. These data strongly suggest that the continuous synthesis of (active) RhoB contributes to the loss of endothelial integrity which is consequent to the inhibition of the ubiquitination cascade. In contrast, RhoA levels and activity are not altered by inhibiting protein synthesis during E1 ligase inhibition, which is in line with the much longer half-life of RhoA (Figure S3A). Thus, these data suggest that regulation of RhoA protein synthesis does not play a major role in control of quiescent endothelial integrity.

Although RhoA and RhoB share high sequence homology and downstream signaling pathways, our data suggest that protein synthesis and ubiquitination play a strikingly different role in regulating their output. It is well established that RhoA activation mediates transient responses such as the rapid (at a time scale of minutes) contraction and relaxation induced by activation of GPCRs such as PAR1, S1PR and LPA receptor (Figure 5E) (Pronk et al. 2019; Yu and Brown 2015). The cycling between the active and inactive form, regulated by RhoGDI, GEFs and GAPs, allows acute, fast (<1 h time scale) and transient activation of RhoA (Figure 6A). The available pool of cytosolic, RhoGDI-bound, inactive RhoA allows these quick responses to receptor agonists, which regulate endothelial integrity. In line with this, our data indicate that ubiquitin-mediated degradation and protein synthesis do not significantly contribute to regulate RhoA activity in quiescent endothelium.

**Figure 6.**
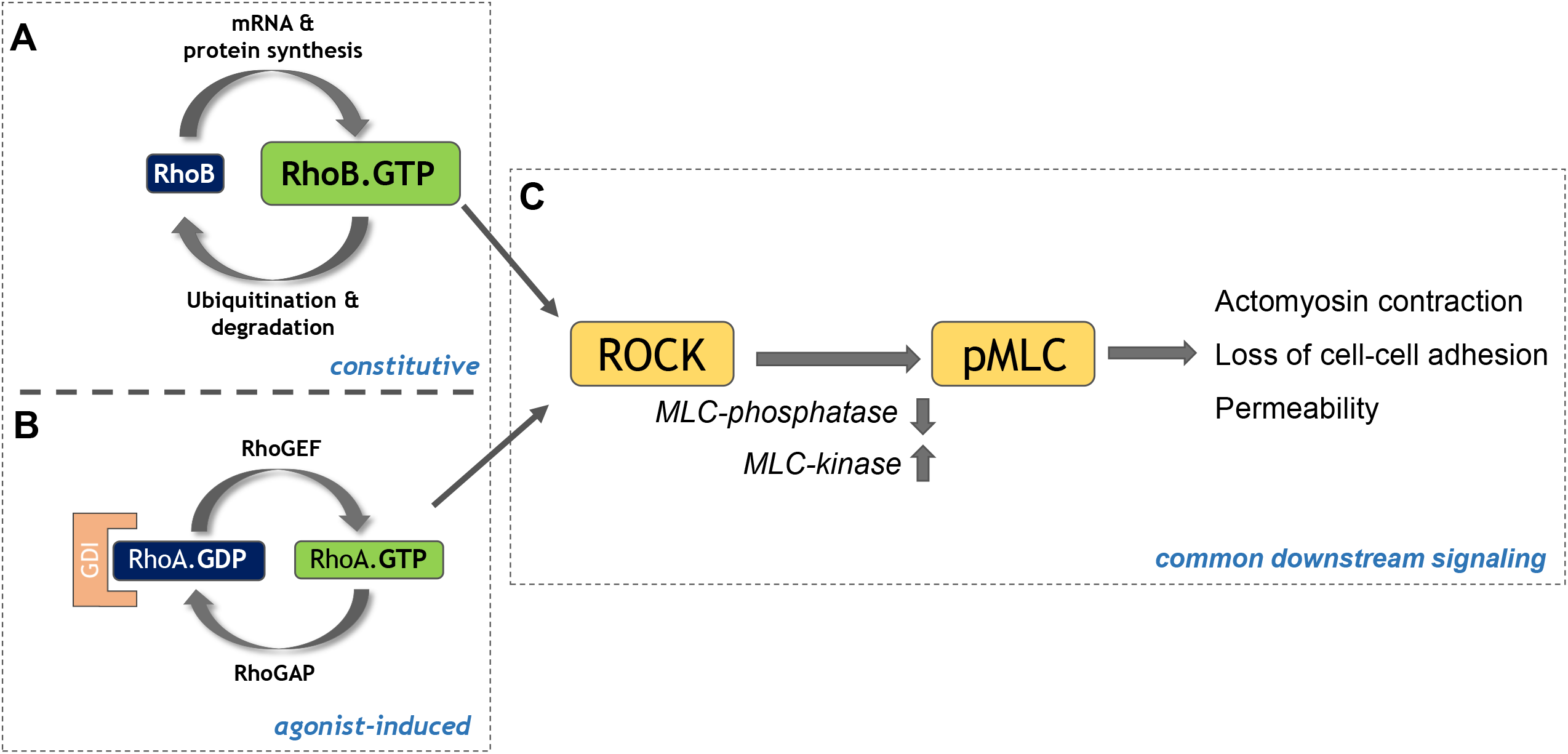
Model of differential regulation of endothelial integrity by RhoB and RhoA. (**A**) The activity of RhoB is mainly regulated at the levels of its mRNA expression, protein synthesis and ubiquitination-mediated degradation, which allows for a relatively stable level of constitutive signaling by RhoB. (**B**) In contrast, GDI, GEF and GAP proteins are primarily responsible to control RhoA activity conveying rapid responses to, mostly GPCR-binding, agonists. (**C**) Despite being differentially regulated, active RhoB and RhoA both signal towards their common effectors, including ROCK, driving the phosphorylation of MLC, which results in actomyosin contraction and an increase in endothelial permeability.

In contrast, the lack of RhoGDI binding and abundance of GTP leads to rapid activation of newly synthesized RhoB. RhoB activity is thus more clearly regulated by a synthesis-degradation cycle (> 2-4 h time scale; Figure 6B). By this mode of regulation, RhoB expression is largely concomitant with its activation and downstream signaling. In contrast to RhoA, this allows RhoB to mediate long term (>hours) and chronic effects. For example, RhoB levels and activity are increased for prolonged periods of time (time scale of hours) due to hypoxia and stimulation by inflammatory mediators such as TNF, IL-1β and lipopolysaccharide (Figure 5A) (Wojciak-Stothard et al. 2012; Marcos-Ramiro et al. 2016; Kroon et al. 2013; Liu et al. 2017). Besides, high RhoB expression was detected in intestinal, inflamed blood vessels of patients with Crohn’s disease (Marcos-Ramiro et al. 2016). Elevated RhoB levels have also been found in urine of patients with chronic kidney disease and RhoB has been identified as upregulated gene 2 - 24h after renal ischemia-reperfusion injury after kidney transplantation (Konvalinka et al. 2016; Zhang et al. 2021). However, once activated, RhoA and RhoB share overlapping downstream signaling, comprising activation of ROCK and the induction of actomyosin-based contraction (Figure 6C).

These findings propose only a limited role for regulation of RhoB activity by GEF and GAP activity in quiescent endothelium, deviating from the prevailing model of Rho GTPase regulation. Recently, Pierce et al. proposed p190BRhoGAP to selectively inactivate RhoB in the context of capillary leak syndrome (Pierce et al. 2017). This could suggest that, under certain conditions or in specific organs, constitutive regulation by GAP proteins may also contribute to RhoB output. Conversely, several studies have identified various E3 ligases targeting RhoA for degradation. For example a role for Cullin3-based ubiquitin ligases targeting RhoA, both in smooth muscle cells as well as in neurons, has been described (Wu, McCormick, and Sigmund 2020; Escamilla et al. 2017). Apparently, under these conditions, GAP-dependent inactivation is insufficient to limit RhoA signaling, and ubiquitin-mediated degradation is prevalent. This may also relate to the fact that in these studies the biological effects (such as blood pressure regulation or synaptic transmission) are of a more chronic rather than an acute nature.

Intriguingly, recent studies described transient RhoB activation upon thrombin or sphingosine-1-phosphate stimulation (Reinhard et al. 2017; Marcos-Ramiro et al. 2016). However, total RhoB protein levels remain unaffected, suggesting that under certain conditions, RhoB acts along with RhoA to mediate rapid responses to GPCRs. Whether there exists a small pool of inactive RhoB available for rapid and transient activation by a GEF in order to boost RhoA-induced signaling, remains to be determined. Single RhoA/B/C depletion induces upregulation of some of the other Rho GTPases (Pronk et al. 2019), indicating possible functional compensation among Rho GTPases.

While we here identify RhoB as part of a proteostatic mechanism that preserves endothelial integrity, there are likely several other proteins, with a similar short half-life that are also involved in controlling endothelial integrity (e.g. c-Jun (Hyer et al. 2018; Fahmy et al. 2006)). Future proteomic analyses will be required to identify such proteins and their associated regulatory mechanisms. This is important, as knowledge on the regulation of endothelial integrity will contribute to our options to target dysregulation of vascular permeability in (inflammatory) disease.

## Acknowledgments

FP was supported by NWO grant OCENW.klein.021

## Author Contributions

P.L.H. conceived the project. P.L.H., F.P. and J.M. designed experiments. F.P., R.W., M.C.O., J.M. conducted experiments and analyzed data. L.A. conducted experiments. F.P. wrote the manuscript. P.L.H. and F.P. edited the manuscript.

## Materials and Methods

### Antibodies and Reagents

The following primary antibodies were used in this study: anti-RhoB (#sc-180, Santa Cruz Biotechnology and #63876, Cell Signaling Technology), anti-VE-cadherin (#2500, Cell Signaling Technology), anti-RhoA (#2117, Cell Signaling Technology), anti-Rac1 (#610650, BD Transduction Laboratories), anit-peIF2α (#9721 Cell Signaling Technology), anti-caspase 9 (#9502, Cell Signaling Technology), anti-cleaved caspase 3 (#9661, Cell Signaling Technology), anti-GAPDH (#2118, Cell Signaling Technology), anti-β-tubulin (#2128, Cell Signaling Technology), anti-Vinculin (#V9131, Sigma-Aldrich), anti-pMLC (#3671, Cell Signaling Technology), anti-ubiquitin (FK2; #A-106, Boston Biochem and #BML-PW8810, Enzo Life Sciences), anti-ICAM1 (#sc-8439, Santa Cruz Biotechnology). As secondary antibodies for western blotting, horseradish peroxidase (HRP)-conjugated goat anti-rabbit and anti-mouse antibodies (Dako) were used.

For immunofluorescent staining the following primary antibodies were used: anti-RhoB (#sc-8048, Santa Cruz Biotechnology) and anti-VE-cadherin (#2500, Cell Signaling Technology). Alexa-488 donkey anti-rabbit and Alexa-555 donkey anti-mouse (Invitrogen) were used as secondary antibodies. The nucleus was stained with DAPI (#62248, Thermo Fisher Scientific) and F-actin with Acti-stain 670 phalloidin (#PHDN1-A, Cytoskeleton).

In this study the inhibitors MLN7243 (#8341, Selleck Chemicals), MLN4924 (#S7109, Selleck Chemicals), MG132 (#S2619, Selleck Chemicals), Y27632 (#Y0503, Sigma), Tunicamycin (#5.04570, Sigma), Cycloheximide (#239764, Merck Millipore Calbiochem), the cytokine TNFα (#300-01A, Peprotech), and the thrombin receptor activator TFLLR-NH2 (trifluoroacetate salt, #T7830, Sigma) were used.

### Cell culture

Human Umbilical Vein Endothelial Cells (HUVEC) were purchased from Lonza (#CC-2519) and cultured on fibronectin-coated plates with Endothelial Cell Medium (ScienCell Research Laboratories). The medium was refreshed every second day. The cells were cultured at 37°C in 5% CO_2_ and used for experiments until passage 5.

Primary human microvascular endothelial cells (hMVEC) were isolated from the foreskin of healthy donors. Informed consents were obtained from all donors in accordance with the institutional guidelines and the Declaration of Helsinki. The cells were isolated as described before (Koolwijk et al. 1996). The primary hMVEC were cultured in M199 medium supplemented with 100□U/mL penicillin and 100□μg/mL streptomycin, 2 mM L-glutamine (all Bio Whittaker/Lonza), 10% heat-inactivated human serum (Invitrogen), 10% heat-inactivated new-born calf serum (Gibco), 150□μg/mL crude endothelial cell growth factor (prepared from bovine brains) and 5□U/mL heparin (Leo pharmaceutical products). The cells were cultured at 37°C in 5% CO_2_ and medium was refreshed every second day. For experiments cells from single donors or pools of 3 donors in passages 5-6 were used.

### siRNA transfection

HUVEC were seeded on fibronectin-coated Electric Cell-substrate Impedance Sensing (ECIS) or culture plates. When cells reached 70-80% confluency siRNA transfection using Dharmafect reagent 1 (#T-2001, Dharmacon) in OptiMEM (gibco) was performed. For gene silencing a final concentration of 25 nM of ON-TARGETplus Human RhoA siRNA SMART pool (siRhoA) or ON-TARGETplus Human RhoB siRNA SMART pool (siRhoB) or combination of both (siRhoA/siRhoB, all Dharmacon) was used. ON-TARGET plus Non-targeting Control pool (N.T.) was used as negative control. After 7 h the transfection medium was replaced by normal culture medium. 72 h post-transfected cells were used for experiments.

### Endothelial barrier function assays

To measure endothelial barrier function, ECIS was performed. HUVEC were seeded on fibronectin-coated ECIS plates containing gold intercalated electrodes (Applied Biophysics). hMVEC were seeded on gelatin-coated ECIS plates. The resistance was monitored at 4000 Hz. When cells formed a stable monolayer, treatment with the compounds was performed as indicated.

Permeability of the endothelial monolayer was measured by passage of horseradish peroxidase (HRP). HUVEC were seeded on gelatin-fibronectin coated Thin-Certs™ (Greiner Bio-One), which have a pore size of 3μm. After forming a stable barrier, HUVEC were treated as indicated. For the last hour of treatment 5μg/ml HRP was added to the upper compartment. After the treatment a sample was taken from the lower compartment and upper compartment (total HRP). The HRP concentration from the lower compartment was calculated as % of the total HRP after adding the substrate tetramethylbenzidine (TMB; Organon Teknika), stopping the reaction with 2M sulfuric acid and by measuring the absorbance at 450nm with a microplate spectrophotometer (Epoch BioTek).

### Western blot

After treatment with compounds as indicated, cells were washed with PBS supplemented with 1 mM CaCl_2_ and 0.5 mM MgCl_2_ and lysed in 2x SDS sample buffer (125□mM Tris-HCl pH 6.8, 4% SDS, 20% glycerol, 100□mM DTT, 0.02% Bromophenol Blue in MilliQ). Proteins were separated by SDS-PAGE and transferred to a nitrocellulose membrane. Membranes were blocked in 5% BSA in TBS-T for 1 h and incubated with designated primary antibodies in 5% BSA in TBS-T overnight at 4°C. After incubation with secondary antibodies, proteins were visualized using enhanced chemiluminescence (Amersham/GE-healthcare) and an AI600 imager (Amersham/GE-healthcare). Densitometric analysis of the band intensities was perfomed using ImageQuantTL.

### Immunofluorescence analysis

HUVEC were seeded on fibronectin-coated 13 mm coverslips (#0117530, Marienfeld superior) and when confluency was reached treated as indicated. After washing twice with PBS supplemented with 1 mM CaCl_2_ and 0.5 mM MgCl_2_, cells were fixed with warm 4% paraformaldehyde in PBS at room temperature for 15 minutes. After three washes with PBS, cells were permeabilized with 0.2% triton X-100 in PBS for 3 minutes and blocked for 1 h with 1% human serum albumin (HSA) in PBS. Hereafter, coverslips were incubated with primary antibodies in 1% HSA in PBS for 1 h at room temperature or overnight at 4°C. Coverslips were washed three times with PBS and incubated with a secondary antibody, Acti-stain 670 phalloidin (#PHDN1-A, Cytoskeleton) or DAPI (#62248, Thermo Fisher Scientific) for 1 h at room temperature. Subsequently, coverslips were washed with PBS and mounted on Mowiol4-88/DABCO solution (Calbiochem, Sigma Aldrich). Confocal scanning laser microscopy was performed on a Nikon A1R confocal microscope (Nikon). Images were analyzed and equally adjusted with ImageJ.

### Rac1 and Rho GTPase activation assays

To analyze Rho activity, confluent HUVEC seeded on fibronectin-coated 55 cm^2^ dishes were treated as indicated. Subsequently, cells were washed with ice-cold PBS supplemented with 1 mM CaCl_2_ and 0.5 mM MgCl_2_ and lysed on ice. The levels of RhoA and RhoB activity were measured using the Rho Activation Assay Biochem Kit (Cytoskeleton) with Rhotekin pulldown according to the manufacturer’s protocol. The input and pulldown samples were analyzed by Western blot.

Rac1 activity was analyzed by performing a CRIB pulldown (Price et al. 2003). HUVEC were seeded on fibronectin-coated 21m^2^ dishes and treated when confluent. Cells were washed with PBS supplemented with 1 mM CaCl_2_ and 0.5 mM MgCl_2_ and lysed in 500 μl ice-cold lysis buffer (150 mM NaCl, 50 mM Tris pH 7.6, 1% Triton-X 100, 20 mM MgCl_2_) containing 30μg biotinylated CRIB peptide. After centrifugation of lysates at 14000rpm and 4°C for 5 minutes, 10% of supernatant was taken as input and mixed with 3x SDS sample buffer (125□mM Tris-HCl pH 6.8, 4% SDS, 20% glycerol, 100□mM DTT, 0.02% Bromophenol Blue). The remaining lysate was incubated with streptavidin beads (Sigma) for 30 min rotating at 4°C. Subsequently, the beads were washed 5 times with lysis buffer with freshly added 10 mM MgCl_2_. The buffer was aspirated and the beads were lysed in 30 μl 2x SDS sample buffer. The input and pulldown samples were analyzed by Western blot.

### Endogenous RhoB immunoprecipitation (IP)

Confluent monolayers of HUVEC grown on fibronectin-coated 55 cm^2^ dishes were treated as indicated. Additionally, all samples were treated with 5 μM MG132 (#S2619, Selleckchem) for 2 h. Cells were washed with PBS supplemented with 1mM CaCl_2_ and 0.5mM MgCl_2_ and lysed in 400 μl ice-cold lysis buffer (50 mM Tris, pH 7.4, 150 mM NaCl, 1 mM EDTA, 1% NP-40 and 1x protease inhibitor cocktail). Lysates were centrifuged at 14000rpm and 4°C for 10 min, and 8% of supernatant was taken as input and mixed with 3x SDS sample buffer (125□mM Tris-HCl pH 6.8, 4% SDS, 20% glycerol, 100□mM DTT, 0.02% Bromophenol Blue). The remaining supernatant was incubated with 1 μg anti-RhoB (#sc-8048, Santa Cruz Biotechnology) and rotated overnight at 4°C. 25 μl Dynabeads Protein G (#10003D, Thermo Fisher Scientific) were added to the lysate and incubated for 1 h rotating at 4°C. The beads were washed four times with lysis buffer and ultimately lysed in 30 μl 2x SDS samples buffer. The input and IP samples were analyzed by Western blot.

### Statistical Analysis

Data are presented as mean ± SD or mean + SD. Statistical analysis was performed using GraphPad Prism. One way ANOVA with Dunnett’s post-hoc test was applied when groups were compared to one (control) group. For comparison of several groups one way ANOVA with Tukey’s post-hoc test was used. p-values <0.05 were considered as statistically significant.

