## Supplemental Figures for "Rapid proteostasis controls monolayer integrity of quiescent endothelium"

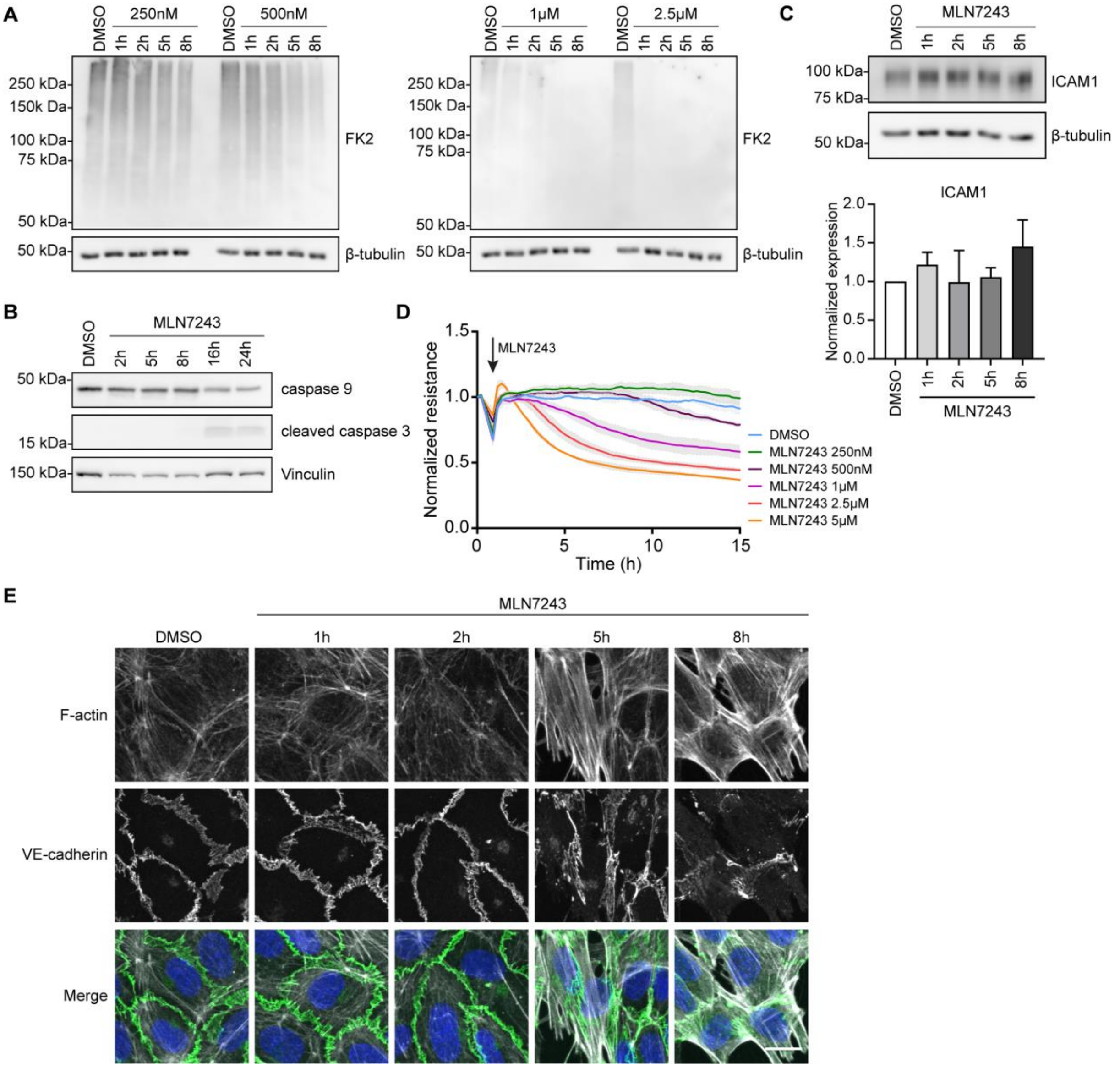


**Figure S1**

(**A**) Western blot analysis for total ubiquitinated proteins (FK2) of HUVEC treated with indicated concentrations of MLN7243 for the indicated times. β-tubulin was used as loading control. (**B, C**) Western blot analysis of HUVEC after treatment with 500 nM MLN7243 for (**B**) ICAM1 and (**C**) caspase 9 and cleaved caspase 3. β-tubulin and vinculin were used as loading controls. Bar graph shows quantification of ICAM1 expression normalized to β-tubulin and DMSO control (data are presented as mean + SD, n=3) (**D**) Normalized resistance of hMVEC treated with different concentrations of MLN7243. (**E**) Zoomed images from Fig 1C: Immunofluorescent staining for F-actin (white) and VE-cadherin (green) and counterstained with DAPI (blue) of HUVEC after treatment with 500 nM MLN7243 for the indicated times. Scale bar represents 15 μm.

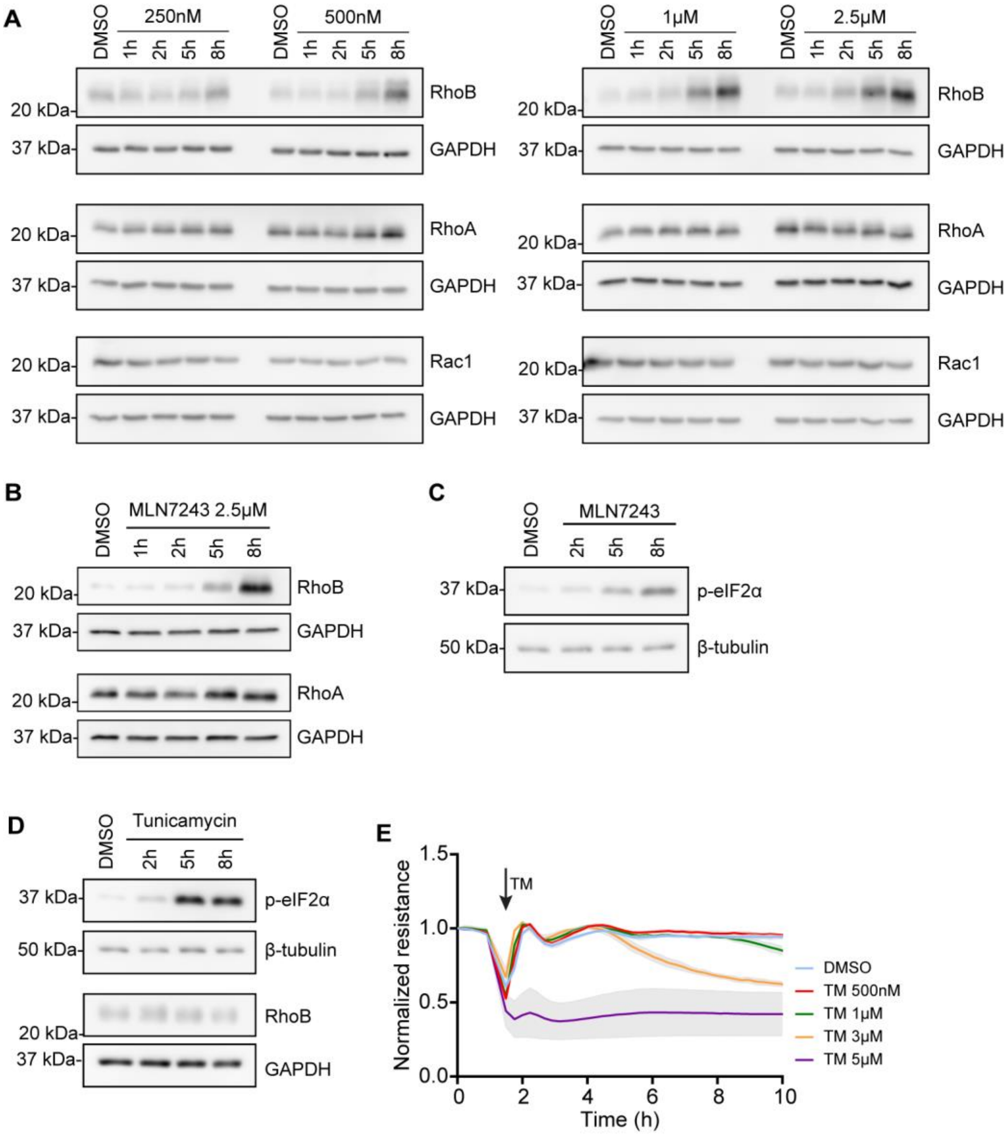


**Figure S2**

(**A**) Western blot analysis of HUVEC after treatment with different concentrations of MLN7243 for indicated times for RhoA, RhoB and Rac1 expression. GAPDH is used as loading control. (**B**) Western blot analysis of hMVEC after treatment with 2.5 µM MLN7243. GAPDH was used as loading control. (**C**) Western blot analysis for expression of p-eIF2α after 500 nM MLN7243 treatment of HUVEC. β-tubulin was used as loading control. (**D**) After treatment of HUVEC with 3 µM Tunicamycin (TM) for indicated times, western blot analysis for expression of p-eIF2α and RhoB. β-tubulin and GAPDH were used as loading controls. (**E**) Normalized endothelial resistance of HUVEC monolayers, treated with the indicated concentrations of Tunicamycin.

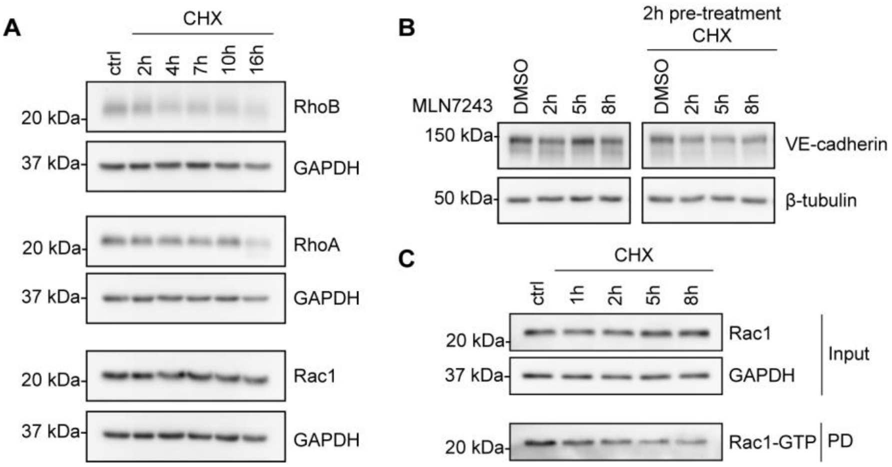


**Figure S3**

(**A, B**) Western blot analysis for (**A**) RhoA, RhoB and Rac1 following treatment as indicated with 0.5 µg/ml CHX and for (**B**) VE-cadherin after 2 h pre-treatment with 0.5 µg/ml CHX and treatment as indicated with 500 nM MLN7243. GAPDH and β-tubulin were used as loading controls, respectively. (**C**) CRIB pulldown from HUVEC treated with 0.5 µg/ml CHX followed by Western blot analysis to detect active Rac1. GAPDH was used as loading control.
